# Inorganic phosphate content does not affect oviposition preference in the invasive pest *Drosophila suzukii*

**DOI:** 10.1101/2022.09.09.507340

**Authors:** Laure Olazcuaga, Robin Guilhot, Jean-Loup Claret, Nicolas O. Rode

## Abstract

The broad variation in host use among polyphagous insects is well documented but still poorly understood. In numerous pest insects, the proximate mechanisms responsible for variation in oviposition preference among host plants remain to be elucidated. The invasive crop pest, *Drosophila suzukii*, attacks a wide range of host fruits. Females prefer ovipositing on particular fruit media (blackberry, cherry, blackcurrant) that are rich in phosphorus. As phosphorus is known to be involved in female reproduction in insect species such as Drosophila, it could drive oviposition preference in *D. suzukii*. Phosphorus is either present as inorganic or organic phosphate in fruits. As the absolute content in macromolecules associated with phosphate in fruits (i.e. proteins and carbohydrates) do not affect oviposition in *D. suzukii*, we tested for the effect of inorganic phosphate on oviposition preference. We measured the egg-laying preferences of *D. suzukii* in a choice environment containing 12 artificial media with increasing content in inorganic phosphate (monopotassium dihydrogen phosphate). In our assay, *D. suzukii* females did not prefer ovipositing in media with high inorganic phosphate content compared to media with lower inorganic phosphate content. As a confirmation, we verified the previous result of a higher female preference for media made of phosphorus-rich fruits (blackberry, cherry, blackcurrant). The higher preference for phosphorus-rich fruits could be driven by macromolecules containing phosphorus (e.g. phospholipids) or by the presence of one or more molecules that do not contain phosphorus, but that happen to be correlated to fruit phosphorus content. Studying the proximate mechanisms driving host use will ultimately help improve the management of *D. suzukii* and other crop pests.

## Introduction

Polyphagous insect pests that infest many host plants are responsible for important economic losses in agriculture (Oerke, 2006). Hence, elucidating the mechanisms that drive patterns of host use by polyphagous insects is an important goal in ecological entomology. Oviposition preference, one of the main traits that determines host use in polyphagous insects, varies widely among host fruits and among varieties within a given host fruit (Singer, 1986). Both the physical (skin thickness, fruit shape, and color) and chemical (nutritional quality, presence of toxins) properties of fruits are thought to affect female oviposition preference (Renwick, 1989).

*Drosophila suzukii* is an insect pest, native to Asia, which invaded both Europe and North America in 2008 (Fraimout *et al*., 2017). This extremely polyphagous pest can infest fruits from at least 19 different plant families (Kenis *et al*., 2016). In particular, it infests soft fruits (strawberry, raspberry, blueberry, mulberry), stone fruits (cherry) and grape, which results in important economic losses for farmers (for example $US 36.1 million in revenue loss between 2009 and 2011 for Californian raspberry producers; Farnsworth *et al*., 2017). Current control methods rely heavily on the use of broad-spectrum insecticides (Schetelig *et al*., 2018). However, as the timing of infestation coincides with ripening, pesticide use is restricted to protect consumers from exposure to chemical residues (Haviland & Beers, 2012). Given the known negative impacts of insecticides on human health and on the environment (Bourguet & Guillemaud, 2016), alternative pest management strategies are deeply needed. Notably, the development of push-pull strategies (e.g., trap-plants) requires a good understanding of the factors that drive oviposition preference in insect pests (Alkema *et al*., 2019).

The roles of certain physical (e.g., Lee *et al*., 2011; Burrack *et al*., 2013; Takahara & Takahashi, 2017; Little *et al*., 2018, 2019; Tait *et al*., 2020) and chemical characteristics (e.g., Dweck *et al*., 2016; Karageorgi *et al*., 2017; Silva-Soares *et al*., 2017; Young *et al*., 2018; see Olazcuaga *et al*., 2019 for a review) in driving oviposition preference of *D. suzukii* are well known. For instance, oviposition preference for a given fruit increases as pH increases and as the force necessary to penetrate the skin decreases (Kinjo *et al*., 2013; Lee *et al*., 2015; Baena *et al*., 2022). The ratio of protein to carbohydrates (P:C ratio) also affects female oviposition preference with females preferring to lay eggs in media rich in carbohydrates relative to proteins (Silva-Soares *et al*., 2017; Young *et al*., 2018). However, the effects of specific molecules of the fruit on oviposition preference have been little investigated (see Olazcuaga *et al*., 2019 for an indirect approach). Olazcuaga *et al*. (2019) found that, in a choice environment, females preferred to lay eggs in media made from fruits rich in phosphorus (specifically, blackberry, cherry, and blackcurrant), resulting in a positive correlation between oviposition preference and phosphorus content within those fruits. Although the existence of the positive correlation between oviposition preference and phosphorus content is not direct evidence of a causal link between phosphorus and preference, it suggests that the presence of large amounts of phosphorus might drive female oviposition preference in *D. suzukii*. Indeed, phosphorus is known to be important for female survival and reproduction in insects, including some Drosophila species (King & Wilson, 1955; Markow *et al*., 1999; Bergwitz, 2012). In both frugivorous and cactophilic Drosophila, the elemental composition of females is richer in phosphorus than that of males (Markow et al., 1999). The authors assumed that females may actively seek phosphorus-rich resources for feeding, as they might use phosphorus for RNA transcription during oogenesis. We thus hypothesize that *D. suzukii* females could seek phosphorus-rich fruits for feeding and potentially for oviposition, resulting in a positive correlation between oviposition preference and the phosphorus content of fruits.

In plant organs such as fruits, leaves, seeds, or roots, phosphorus is mostly present as inorganic phosphate, usually associated with potassium, or as organically bound phosphate (e.g. sugar phosphates, phospholipids, nucleotides; Lott *et al*., 2000; Marschner, 2012). Organically bound phosphate is unlikely to affect oviposition preference, as the absolute content of macromolecules associated with phosphate in fruits such carbohydrates or proteins do not affect oviposition preference in *D. suzukii* (Olazcuaga *et al*., 2019). We hypothetized that female oviposition might be driven by the inorganic phosphate content of fruits and tested whether oviposition preference of *Drosophila suzukii* does depend on inorganic phosphate content.

## Material and methods

### Fly population and maintenance

To minimize phenotypic variation, we used an inbred line of *D. suzukii* (WT3 2.0; Paris *et al*., 2020). Inbred lines represent powerful tools to unravel potential drivers of oviposition preference of natural populations of *D. suzukii* (Karageorgi *et al*., 2017). Individuals were maintained in vials with standard laboratory fly food (GF, Backhaus *et al*., 1984) for several discrete generations in a growth chamber (15 days of larval development, 6 days of adult maturation and 24h to lay eggs; 21°C (+/- 2°C), 65% relative humidity and 16:8 (L:D) light cycle). We performed oviposition preference assays under the same environmental conditions.

### Preparation of media with increasing content in inorganic phosphate

To test whether the oviposition preference of *D. suzukii* depends on the inorganic phosphate content, we supplemented a minimum fly food medium with increasing amounts of monopotassium dihydrogen phosphate (KH_2_PO_4_). Other sources of phosphorus (i.e. calcium or iron phosphate) are less likely to be used by plants due to their insolubility (Lindsay C. Campbell, pers. com.). The minimum medium was made by adding 1% (w/v) of agar, 6% of inactive malted brewer’s yeast, 1.5% of yeast extract, and an antimicrobial solution (consisting of 6 ml of 99% propionic acid, 10 ml of 96% ethanol and 0.1% of methylparaben sodium salt) to 1000 ml of sterile deionized water (Table S1). Since KH_2_PO_4_ is a good buffer, we adjusted the pH of each medium to 4 (i.e., the pH of fruit media used in Olazcuaga et al. 2019 and in the present assay) using hydrochloric acid (HCl).

To maximize the ability to detect an effect of the inorganic phosphate content on oviposition preference, we varied the inorganic phosphate content from 0 to 4.7 g/kg medium. This range corresponds to a monopotassium dihydrogen phosphate ranging from 0 to 6.6 g/kg medium and phosphorus content ranging from 0 to 1.5 g/kg medium, several times larger than the range of phosphorus content observed in the 12 artificial fruit media used in Olazcuaga *et al*., 2019 (0 to 0.30 g/kg medium).

### Standardization of the age of the flies

Following Olazcuaga et al. 2019, we used groups of 6-day old flies for the oviposition experiment (i.e., sexually mature females; Emiljanowicz *et al*., 2014). The artificial media used for the experiment were freshly made and flies were not previously exposed to fruit media, two factors known to influence oviposition site selection in *D. suzukii* (Hoffmann, 1985; Tait *et al*., 2020; Elsensohn *et al*., 2021b, 2021a). To maximize the number of replicates and minimize the stress of handling adults with CO2 to control sex ratio, we did not control or measure the sex ratio of the adults in each group of 20 individuals. As we used a very large number of replicates (see below), our oviposition preference assay represents that of a population with an average sex ratio of 50:50.

### Oviposition preference assay

Following Olazcuaga *et al*., 2019, oviposition preference was measured in oviposition arenas (choice assay) with 12 compartments. We randomly distributed each of 12 different inorganic phosphate content media across the 12 compartments of each oviposition arena. In each arena, we placed a group of 20 flies aged 6 days, as described above. After 24h, we counted the number of eggs laid in each compartment. This method is a direct approach to measuring phosphate preference and is, therefore, more powerful than other types of indirect approaches (e.g., measuring preference in arenas containing 12 puree fruits where the phosphate content would have been equalized among the fruits), thus avoiding potential false negative results. We replicated this assay 129 times. In addition, to confirm the existence of a positive correlation between oviposition preference and phosphorus content of fruit media, we performed the same protocol as Olazcuaga *et al*. (2019) using the 12 artificial fruit media that vary in phosphorus content. To estimate the number of replicate arenas for this confirmatory assay (hereafter “fruit assay”), we performed a power analysis with simulations based on the subsampling of the original data (see Sup. Mat.). We found that an assay with 6 fruit arenas provided enough power to detect a significant effect of phosphorus content on oviposition preference in more than 99% of the simulations. Hence, we used 20 fruit arenas for the fruit assay. The position of 129 test arenas and 20 fruit arenas was randomized within the growth chamber.

### Statistical analyses

To investigate whether oviposition preference depends on inorganic phosphate, we tested for the effect of the inorganic phosphate content on the number of eggs laid in each compartment using a Generalized Linear Mixed Model with a log link (Poisson distribution for count data). The model included inorganic phosphate content as a fixed effect (continuous variable). To control for potential correlations in the number of eggs among compartments across and within arenas, we added the position of the compartment within the arena and arena identity as random effects (with 12 and 129 levels respectively). To account for overdispersion, we added an observation-level random effect (1548 levels). We tested for the effect of inorganic phosphate content using a Likelihood Ratio Test (LRT; Bates *et al*., 2015). To quantify the proportion of variance in oviposition preference explained by inorganic phosphate content, we computed the likelihood-based coefficient of determination using the *rr2* package (Ives, 2019). In addition, we separately performed the same analysis on the data from the 20 fruit arenas using the phosphorus content of the 12 fruit media as estimated in Olazcuaga *et al*., 2019.

## Results

### Testing for a causal effect of inorganic phosphate on oviposition preference

The number of eggs laid in each compartment did not increase with inorganic phosphate content (LRT Χ^2^ = 2.94, df = 1, *P*-value = 0.09; Fig. 1A). In contrast, the number of eggs laid in each compartment increased with the phosphorus content of the 12 fruit media in fruit arenas (LRT Χ^2^ = 23.53, df = 1, *P*-value < 0.001; Fig. 1B).

**Fig. 1:**
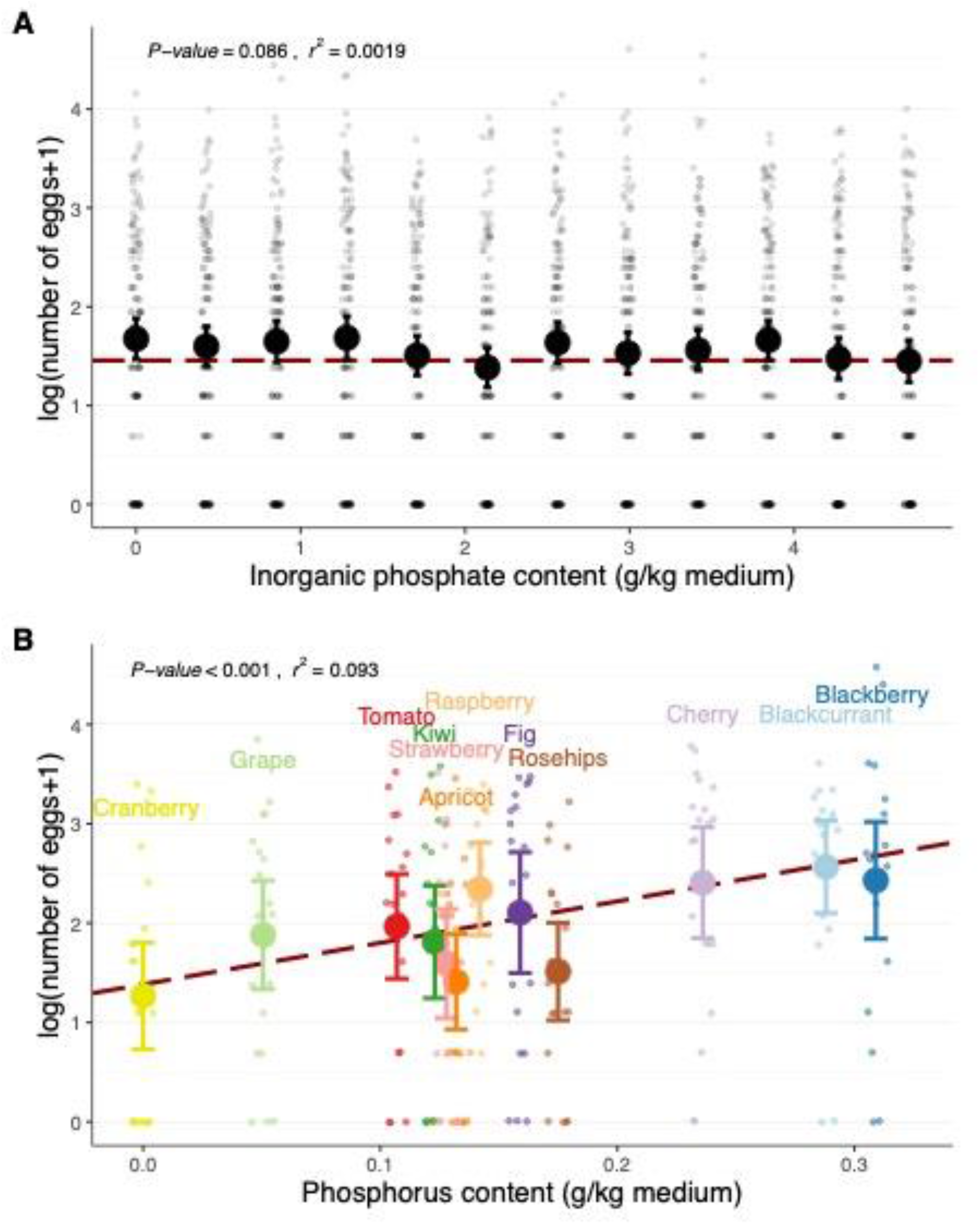
Relationship between female oviposition preference and (A) increasing content in inorganic phosphate supplemented in standard medium or (B) increasing content in phosphorus in 12 different artificial fruit media. Large and dark filled circles with error bars represent the mean with 95% confidence intervals, small and light filled circles represent the observed data. The dotted red line represents the number of eggs predicted by the best model. The *P*-value and coefficient of determination of the effect of phosphate or phosphorus content on oviposition preference are given above each panel.

## Discussion

We used an inbred line of *D. suzukii* to test whether inorganic phosphate content affects oviposition preference in this invasive pest. Our results confirm previous findings of a greater oviposition preference for fruit media with the highest phosphorus content. However, females did not prefer to lay eggs on minimum media with a high inorganic phosphate content.

### Identifying molecules involved in oviposition preference

Two hypotheses may explain why we found females prefer to oviposit in fruit media with high phosphorus content, but not in minimum media with high inorganic phosphate content. First, although the content in macromolecules associated with phosphorus such carbohydrates or proteins do not drive oviposition preference in *D. suzukii* (Olazcuaga *et al*. 2019), other macromolecules (e.g. phospholipids) might drive oviposition preference. Second, oviposition preference could be driven by one or more molecules that contain no phosphorus, but whose content may be correlated with phosphorus content across the 12 fruits studied. Some volatile compounds, such as linalool, are known to be involved in the oviposition choice of *D. suzukii* (Baena *et al*., 2022). However, whether the presence of these compounds correlates with phosphorus content remains to be determined.

### Deciphering mechanisms that may influence oviposition preference

Two proximate mechanisms could explain the oviposition preference for specific fruits (Visser, 1986). First, females might be attracted by olfactory compounds which are honest cues of the quality of the fruit substrate (olfaction-driven preference; Bruce *et al*., 2005). In this scenario, oviposition preference would be an active choice made by females for oviposition (i.e., associated with a non-random flight targeted towards the best fruit substrate for either female feeding or egg development). Second, females might not be attracted by olfactory compounds, but may spend more time on fruits that are more nutritious for them (gustation-driven preference; Amrein & Thorne, 2005). If females randomly lay eggs when they feed, they might eventually lay more eggs on fruits they feed on the most. To dissociate these two mechanisms, using a preference experiment, Clymans *et al*., (2019) showed that *D. suzukii* females are attracted to fermentation volatiles when searching for food and to fruit volatiles when searching for an oviposition substrate. Similar experiments would allow for a better understanding and identification of the factors driving host preference in *D. suzukii*. Finally, the detection of specific volatile compounds could help to improve the design of traps to increase pest trapping efficiency and ultimately improve the prediction of potential damages (Burrack *et al*., 2015).

### Integration of the results in the context of natural populations

Although the experimental conditions differ from natural conditions, especially with the use of an inbred line and fruit purees, the results found in this study represent a strong basis to draw inferences on oviposition preference in natural populations for two reasons. First, fruit media may provide a faithful sensorial representation of fruits in the field. Indeed, we recently found that female oviposition preference was higher on fruit medium corresponding to the fruit from which the female originated than on fruit media corresponding to other fruits (Olazcuaga *et al*., 2022). This discrimination of *D. suzukii* females among different artificial fruit media suggests that these media do harbor the same cues as natural fruits. Second, the behavior of an inbred line accurately represents that of a natural population. Indeed, oviposition preference on 12 artificial fruit media was consistent between the inbred line (this study) and our previous study (Olazcuaga *et al*., 2019).

### Conclusion

By addressing an important and long-standing question regarding the factors that drive oviposition preference in crop pests, our study proposes a straightforward methodology to identify chemical compounds that may determine oviposition preference in an insect pest. This methodology can be used to test for the effects of other chemical compounds on the oviposition preference of important insect pests.

## Acknowledgments

We are very grateful to Graeme D. Batten and Lindsay C. Campbell for insightful discussions on the chemical composition of fruits and Ruth A. Hufbauer for comments on the manuscript and for insightful discussions. We also thank two anonymous reviewers for their useful comments. L.O. acknowledges support from the European Union program FEFER FSE IEJ 2014-2020 (project CPADROL), the INRAE scientific department SPE (AAP-SPE 2016) and the US National Science Foundation (DEB-1930650 to Ruth Hufbauer). N.O.R. acknowledges support from the CeMEB LabEx/University of Montpellier (ANR-10-LABX-04-01).

## Author contributions

Conceptualization: L.O., N.O.R., R.G.

Experimental design and data acquisition: L.O. and J-L.C with inputs of R.G. and N.O.R. Statistical analyses and writing: L.O. and N.O.R with inputs of R.G. and J-L.C.

## Data availability statement

The data and R scripts for our analyses are available at: https://github.com/olazlaure/PhosphorusPreference2022

They will be archived on Figshare upon acceptance of the manuscript.

