## Supplementary materials for "Inorganic phosphate content does not affect oviposition preference in the invasive pest *Drosophila suzukii*"

### Appendix S1: Statistical power to determine the number of fruit arenas

##### *Motivation and methods*

To determine the number of replicates of the fruit arenas needed to detect an effect of fruit phosphorus content on the number of eggs laid, we performed a power analysis based on the oviposition preference data published in Olazcuaga *et al.*, 2019. Briefly, we subsampled without replacement *n* arenas (*n* ranging from 1 to 70) among the 70 arenas in the original dataset. We then used the model presented in the main text to test whether the effect of phosphorus content on the number of eggs was significant using a Likelihood Ratio Test (see main text). We simulated 100 datasets for each value of *n*, the number of subsampled arenas. Our statistical power was estimated as the percentage of simulations where we could detect a significant effect of the fruit phosphorus content on the oviposition preference.

##### *Results*

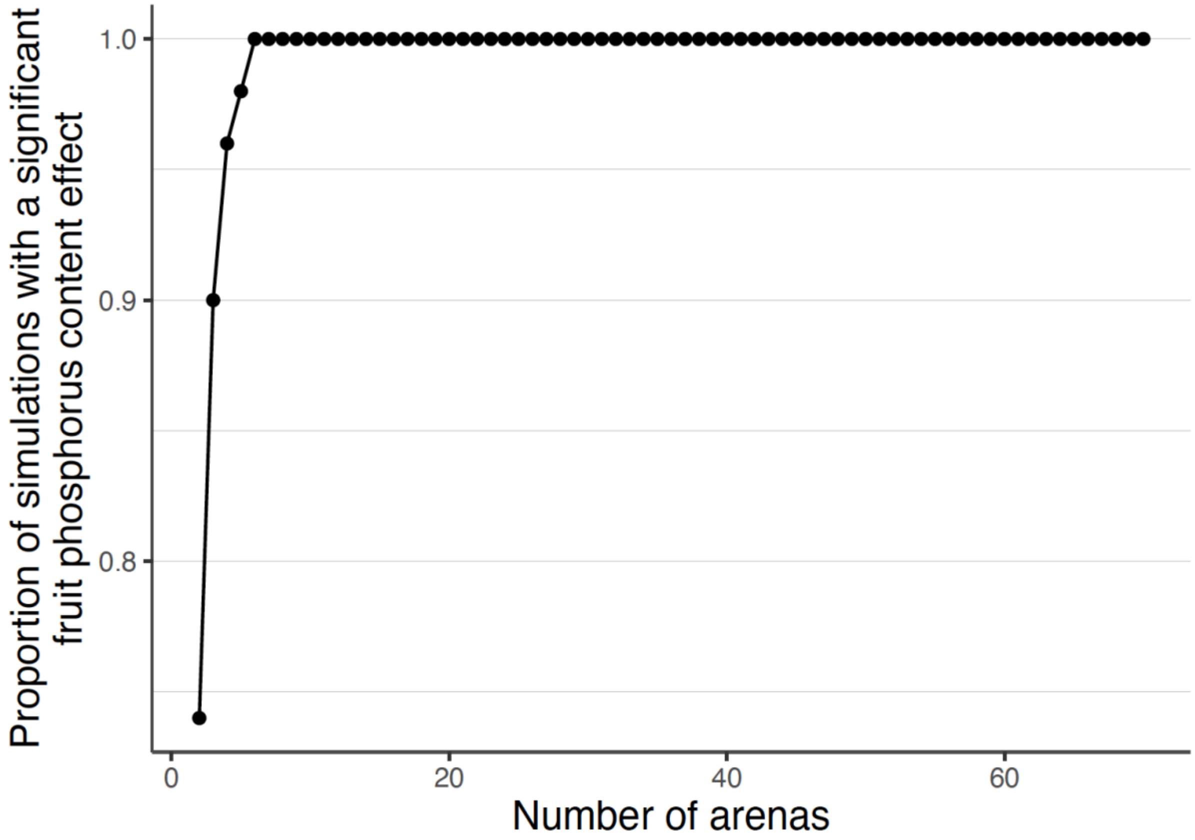


**Figure S1. Estimation of the power of detecting an effect of phosphorus content of fruit medium for an increasing number of subsampled arenas (*n*).**

Statistical power increased with the number of subsampled arenas (*n*). Most importantly, we detected an effect of the fruit phosphorus content with a power greater than an arbitrary threshold of 99% for a number of subsampled arenas higher than 6.

#### *Conclusion*

From the power analysis results given above, we conclude that six control arenas are sufficient to detect a significant effect of the fruit phosphorus content on the oviposition preference.

### Supplementary Table

**Table S1. Recipe and reference of products used to make the minimum medium.**
The density used for the determination of the masses of the liquids was that at room temperature (25 °C). NA = reference not available. * note that a H2PO4 mass of 5.17g is equivalent to a content of 4.70 g/kg medium).

| Product | Mass (g) | Commercial reference | Supplier |
|---|---|---|---|
| Distilled water | 1000 | - | Lab production |
| Agar | 10 | 20768,361 | Vwr |
| Inactive brewer's yeast | 60 | 25108,10,3 | Biocoop |
| Inactive yeast extract | 15 | SI-92144-5KG-F | Analytic lab |
| Alcohol | 7.9 | 78 602 005 | Meridis |
| Methylparaben sodium salt | 1 | SI-H5501-500G | Analytic lab |
| Propionic acid | 5.94 | 8.00605.2500 | Vwr |
| KH2PO4 | [0;…; 7.25] | 795496 | Sigma |
| (equivalence H2PO4) | [0;…; 5.17] | | |
| (equivalence P) | [0;…; 1.5] | | |
